# Aligning protein generative models with experimental fitness via Direct Preference Optimization

**DOI:** 10.1101/2024.05.20.595026

**Authors:** Talal Widatalla, Rafael Rafailov, Brian Hie

## Abstract

Generative models trained on unlabeled protein datasets have demonstrated a remarkable ability to predict some biological functions without any task-specific training data. However, this capability does not extend to all relevant functions and, in many cases, the unsupervised model still underperforms task-specific, supervised baselines. We hypothesize that this is due to a fundamental “alignment gap” in which the rules learned during unsupervised training are not guaranteed to be related to the function of interest. Here, we demonstrate how to provide protein generative models with useful task-specific information without losing the rich, general knowledge learned during pretraining. Using an optimization task called Direct Preference Optimization (DPO), we align a structure-conditioned language model to generate stable protein sequences by encouraging the model to prefer stabilizing over destabilizing variants given a protein backbone structure. Our resulting model, ProteinDPO, is the first structure-conditioned language model preference-optimized to experimental data. ProteinDPO achieves competitive stability prediction and consistently outperforms both unsupervised and finetuned versions of the model. Notably, the aligned model also performs well in domains beyond its training data to enable absolute stability prediction of large proteins and binding affinity prediction of multi-chain complexes, while also enabling single-step stabilization of diverse backbones. These results indicate that ProteinDPO has learned generalizable information from its biophysical alignment data.

## 1. Introduction

Machine learning models for biological structure and function prediction have seen tremendous improvements due to large-scale unsupervised training on databases such as UniProt and the Protein Data Bank (PDB) [28, 24, 2, 57, 17, 5, 48]. One such class of models are structure-informed language models, or “inverse-folding” models, which generate protein sequences that fold into a given structure [17, 20, 9]. These models are surprisingly capable of guiding improvements in binding affinity [45] and improving protein expression, solubility, and stability [52] despite not observing these properties, nor any explicit biophysical information, during training.

These capabilities most likely emerge because a diverse training dataset naturally contains examples that possess these properties. For example, many of the structures in the PDB are of stable or stabilized proteins [12], so models trained on the PDB demonstrate modest zero-shot stability prediction. However, evolutionary selection does not only optimize natural proteins for stability. Consistent with this, unsupervised models are still inferior at stability scoring compared to supervised models such as ThermoMPNN [11]. Using a model trained on unlabeled data for stability-related tasks results in an “alignment gap”: a disagreement between the information learned by the unsupervised model and the specific definition of protein fitness.

A solution would be to explicitly provide the generative model with task-specific information while retaining the model’s generative capabilities. A simple way of doing this is with supervised finetuning (SFT) in which we further train the model on a curated set of positive examples according to a property of interest [33, 43]. However, SFT only maximizes the likelihood over a set of positively labeled samples, making it easy for the model to overfit to the finetuning dataset and lose the general information learned during unsupervised pretraining. More broadly, finding better ways to incorporate experimental fitness information into protein generative models is an important open question.

Direct Preference Optimization (DPO) is an algorithm for lightweight alignment of generative models [39]. DPO enables us to explicitly inform the model of examples that may appear similar at a surface level but are actually very different when considered according to a property of interest (**Figures 1A,B**). This is especially useful in biological settings where single amino-acid changes can have global effects on stability and structure [47]. By providing the model with positive and negative labels of similar proteins, DPO discourages overfitting to a set of positive examples and uses the full fitness landscape to inform the model’s predictions (**Figure 1D**).

**Figure 1.**
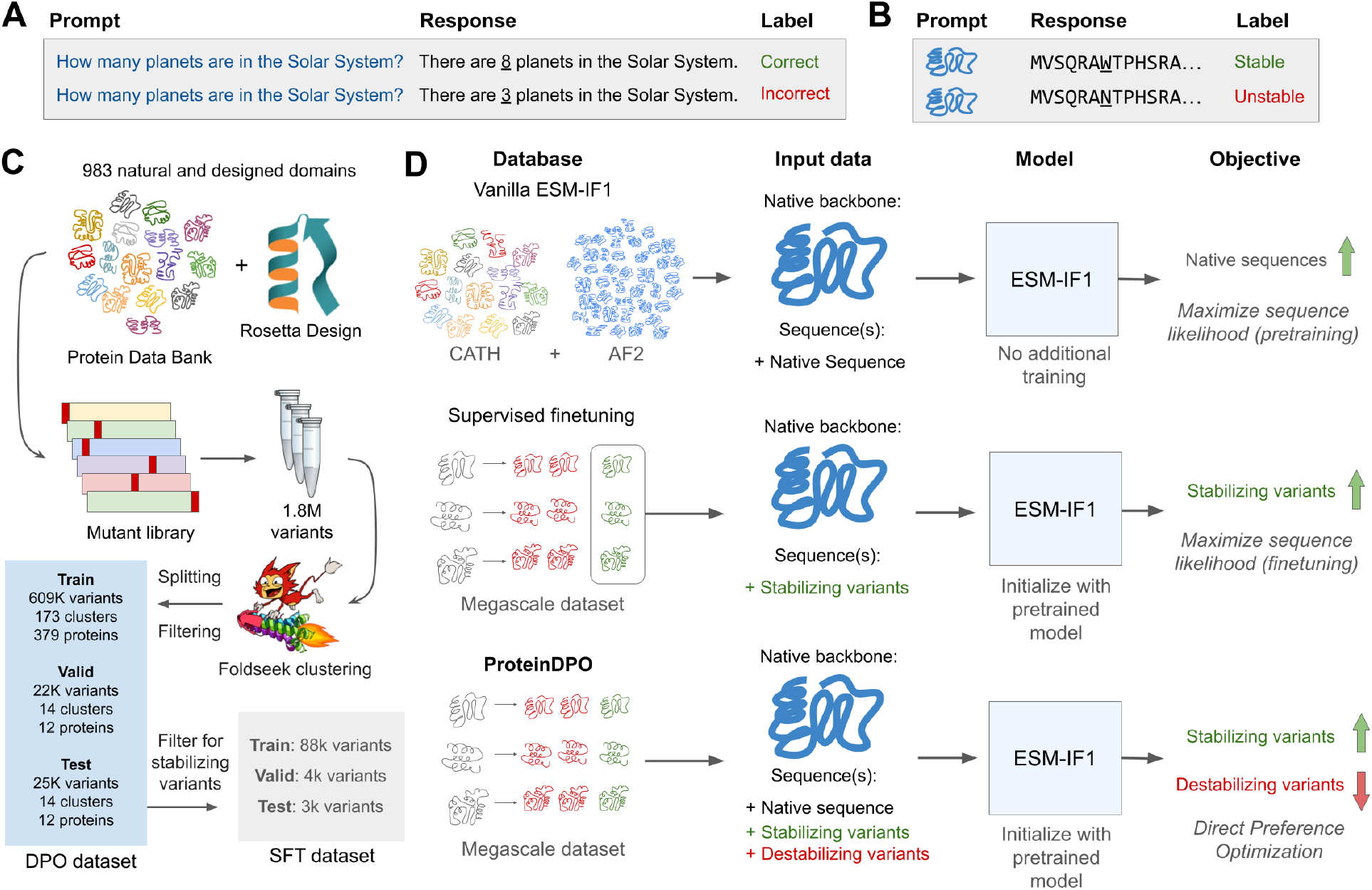
Dataset curation and model training. **(A)** DPO was developed to align the responses of language models to labeled human preferences given a text prompt. **(B)** Analogously, we align a structure-conditioned language model to perform thermostable sequence design using protein backbone conditioning as a “prompt,” sequence generation as a “response,” and experimental fitness as “preference” data. **(C)** Overview of the curation of DPO and supervised finetuning datasets from the Megascale dataset, which contains stability measurements of ∼1.84M sequence variants from 983 protein domains from the PDB and Rosetta design [55, 1, 40]. While filtering out unreliable ΔΔG measurements, we also structurally cluster the protein domains with FoldSeek at a threshold of 0.5 similarity to alleviate leakage between training splits. From these same splits used for DPO, variants with stability superior to the native protein (ΔΔG < 0) were selected for supervised finetuning. **(D)** Overview of the training procedure for the models compared in this study: vanilla ESM-IF1, supervised finetuned ESM-IF1, and ProteinDPO. Vanilla ESM-IF1 was developed and trained by Hsu et al. [17] with maximum likelihood training on structure-conditioned, autoregressive token prediction with protein domains from CATH [25] and AlphaFold2-predicted [24] structures (AF2) [17]. The supervised finetuned model was trained in the same manner, but initialized with the vanilla weights and trained on stabilizing variants from the curated Megascale dataset. ProteinDPO was trained on the curated Megascale on variants of all fitness with a DPO objective (**Methods**).

In this study, we use DPO to imbue stability information into a pretrained structure-conditioned language model, ESM-IF1 [17], using experimental measurements from the “Megascale” stability dataset [55] containing approximately 1.84 million sequence variants across 983 diverse proteins domains (**Figure 1C**). Our aligned model, which we refer to as ProteinDPO (**Protein D**irect **P**reference **O**ptimization), substantially improves upon the stability scoring of vanilla ESM-IF1 and a model trained via supervised finetuning (SFT) on a subset of stable variants from Megascale, and even performs competitively to supervised and physics-based models on single-mutant stability prediction. Despite being trained solely on the relative changes in stability (ΔΔG) of small protein monomers, ProteinDPO also generalizes to scoring absolute stability (ΔG) of larger unrelated proteins, the thermal melting point of antibodies, and the binding affinity of large complexes, which are regimes inaccessible to several supervised models. Beyond scoring, using ProteinDPO to generate new sequences results in designs that have higher stability compared to the native protein and to generations produced by the vanilla and SFT models when evaluated with an *in-silico* force-field. These results establish DPO as an effective method for aligning a generative model with an experimental fitness landscape.

## 2. Results

We repurposed DPO, originally designed to align the *responses* of natural language models to text *prompts* using human *preference* labels [39] (**Figure 1A**), to instead align the generation of protein sequences (“responses”) given their native structure (“prompts”) using their respective experimentally measured stability (“preferences”) (**Figure 1B**). We train and compare the previously described DPO reward models that utilize preference ordered pairs and rankings as input (referred to as Paired and Ranked, respectively). In this study, we also introduce a third, new DPO training objective (Weighted) that directly incorporates scalar labeled data (**Methods**).

### 2.1. Megascale training and relative stability prediction performance

All ProteinDPO models were trained on the Megascale dataset with the DPO objective using native backbones and stability ordered pairs, rankings, or scalar-labeled sets of sequence variants as input, depending on the reward model (**Methods**). To prevent data leakage between training and evaluation splits, we structurally clustered the variants by native structure with FoldSeek [56] and randomly distributed clusters across splits. To determine whether DPO and SFT training improved upon the zero-shot capability of vanilla ESM-IF1 to score changes in stability upon mutation (ΔΔG) (**Figure 2A**), we evaluated the average correlation of mutant sequence likelihoods with ΔΔG across each native protein in the holdout set (**Figure 2B**). While the SFT model modestly improves (Pearson *R* = 0.59, Spearman *ρ* = 0.57) from vanilla ESM-IF1 (Pearson *R* = 0.55, Spearman *ρ* = 0.53), ProteinDPO clearly surpasses both models (Pearson *R* = 0.72-0.73, Spearman *ρ* = 0.69-0.72). We also report the average AUROC, representing the models’ ability to distinguish between stabilizing (ΔΔG < 0) or destabilizing (ΔΔG > 0) variants, and observe the same trend in the vanilla (AUROC = 0.74), SFT (AUROC = 0.76) and ProteinDPO models (AUROC = 0.82-0.84).

**Figure 2.**
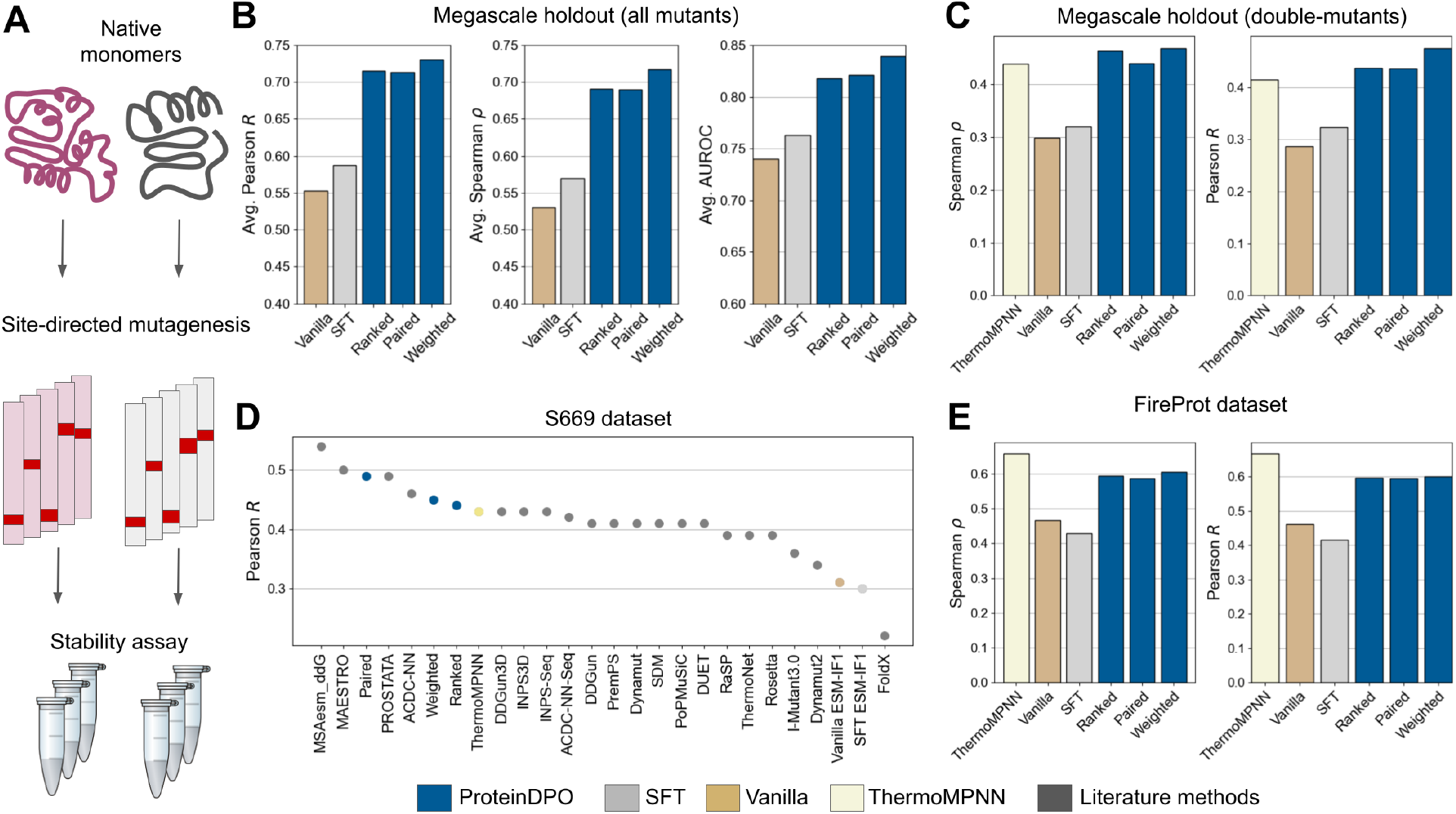
ProteinDPO achieves competitive mutant stability prediction. **(A)** We benchmarked models using deep mutational scanning (DMS) datasets that perform stability measurements over an exhaustive set of variants of protein monomers. Protein stability datasets such as FireProt [51] and S669 [36] are compiled from DMS experiments across diverse monomeric proteins, in which certain residues are mutated, variants are expressed, and their stability is measured experimentally. **(B)** Performance of models on predicting stability changes upon mutation (ΔΔG) in curated Megascale holdout data, including both single and double-mutant variants. We report the correlation of mutant sequence likelihoods with their experimentally measured ΔΔG averaged across each native protein in the holdout data. AUROC measures the models’ ability to classify between variants which increase (ΔΔG < 0) or decrease (ΔΔG > 0) stability compared to native sequence. **(C)** Trained on the same training data as ThermoMPNN, we show the correlation between all model predictions (ThermoMPNN) or likelihoods (Vanilla, SFT, and ProteinDPO) with ΔΔG of all double-mutant variants in ThermoMPNN holdout data [11]. Since ThermoMPNN cannot score multiple mutations, we add the predicted ΔΔGs of the two respective single-mutations as a proxy. **(D)** Performance of ProteinDPO, SFT, vanilla ESM-IF1, and literature methods on the S669 dataset. Performance of literature methods are provided from Cuturello et al. [36, 8] **(E)** Performance of models on predicting single mutant stability changes (ΔΔG) with a version of the FireProt dataset [51] filtered of proteins homologous to those in Megascale [11].

Additionally, we sought to evaluate performance specifically on Megascale variants that have multiple mutations relative to the native sequence. Along with SFT and vanilla ESM-IF1, we compare with ThermoMPNN, a state-of-the-art supervised model based on ProteinMPNN [9] also trained on Megascale. For this comparison, we retrained our models on the training split from Diekhaus et al. [11] and evaluated on the double-mutant variants belonging to their holdout proteins. Notably, ProteinDPO can be trained with double-mutant variants (although we did not for this evaluation for a fair comparison) and can also naturally evaluate the likelihood of combined mutants. In contrast, the architecture and training procedure of ThermoMPNN restricts the model to single mutations. We found that all models do significantly worse (Pearson *R* and Spearman *ρ* = 0.3-0.5) (**Figure 2C**) compared to their ability to score single-mutations (Pearson *R* and Spearman *ρ* = 0.5-0.8) [11], suggesting the non-linear effects of simultaneous mutations, which can energetically influence one another [16], pose a challenge to stability prediction. When we score these variants as a linear combination of single-mutants by adding likelihoods (ProteinDPO) or scores (ThermoMPNN) of only the individual mutations, ProteinDPO underperforms ThermoMPNN (ProteinDPO Pearson *R* = 0.36-0.40, ThermoMPNN Pearson *R* = 0.41) (**Table S1**). However, when we utilize the likelihood summed across the whole sequence with our method, all ProteinDPO models surpass ThermoMPNN (Pearson *R* = 0.44-0.48) (**Figure 2C**), which is fundamentally constrained to the “additive” regime. This suggests ProteinDPO benefits from synthesizing information across the full sequence to capture non-linear effects of multiple mutations.

### 2.2. Comparison with other methods for predicting relative stability

Next, we evaluated if ProteinDPO, trained on Megascale proteins (40 - 72 residues), can generalize to other datasets with larger (up to ∼300 residues) and unseen folds: S669 and FireProt. These datasets are compilations of experimental stability measurements of mutants derived from several deep mutational scanning (DMS) experiments of native proteins (**Figure 1A**) and are popular benchmarks of protein stability predictors. To alleviate data leakage with S669, we retrained ProteinDPO models on a S669-homologue-free Megascale training set from Cutrello et al. [8], and to alleviate leakage with FireProt we evaluated on a Megascale-homologue-free FireProt curation from Diekhaus et al. [11].

We observed that ProteinDPO ranks highly amongst supervised and physics-based ΔΔG predictors (Figure **2D**), some of which were also trained on Megascale. Additionally, we find ProteinDPO is competitive with ThermoMPNN [8, 11] on the respective S669 [36] (ProteinDPO Pearson *R* = 0.44-0.47, ThermoMPNN Pearson *R* = 0.43) (**Figure 2D**) and FireProt [51] (ProteinDPO Pearson *R* = 0.6, ThermoMPNN Pearson *R* = 0.65) (**Figure 2E**) test sets. Interestingly, we find the SFT model, despite improving upon vanilla ESM-IF1 on Megascale holdout prediction (Pearson *R* increase of 0.03, Spearman *ρ* increase of 0.04), does slightly worse on the FireProt (Pearson *R* decrease of 0.05, Spearman *ρ* decrease of 0.04) and S669 datasets (Pearson *R* decrease of 0.01), while the improvement of ProteinDPO over the Vanilla model generalizes to FireProt (Pearson *R* and Spearman *ρ* increases of 0.13-0.14) and S669 (Pearson *R* and Spearman *ρ* increases of 0.13-0.16) as well, suggesting the SFT model has overfit to the training data while ProteinDPO has incorporated generalizable stability information.

### 2.3. Absolute stability prediction

Encouraged by our finding that using whole-sequence likelihoods with our method improves upon addition of single-mutant likelihoods for scoring the *relative* stability (ΔΔG) of double-mutatant variants in Megascale, we sought to evaluate if ProteinDPO can use whole-sequence reasoning to rank *absolute* stability (e.g., ΔG) measurements of both native proteins and antibodies. Antibodies are relevant therapeutic proteins for a wide array of diseases, for which high thermostability is important for developability and storage [4, 41, 58]. To this end, we evaluate on a dataset of 26 natural protein absolute stabilities (ΔG) from Maxwell et al. [31], and thermal melting points (*T*_mid_) of 483 clinical and human derived antibodies from Jain et al. and Shehata et al. [46, 21] (**Figure 3A**).

**Figure 3.**
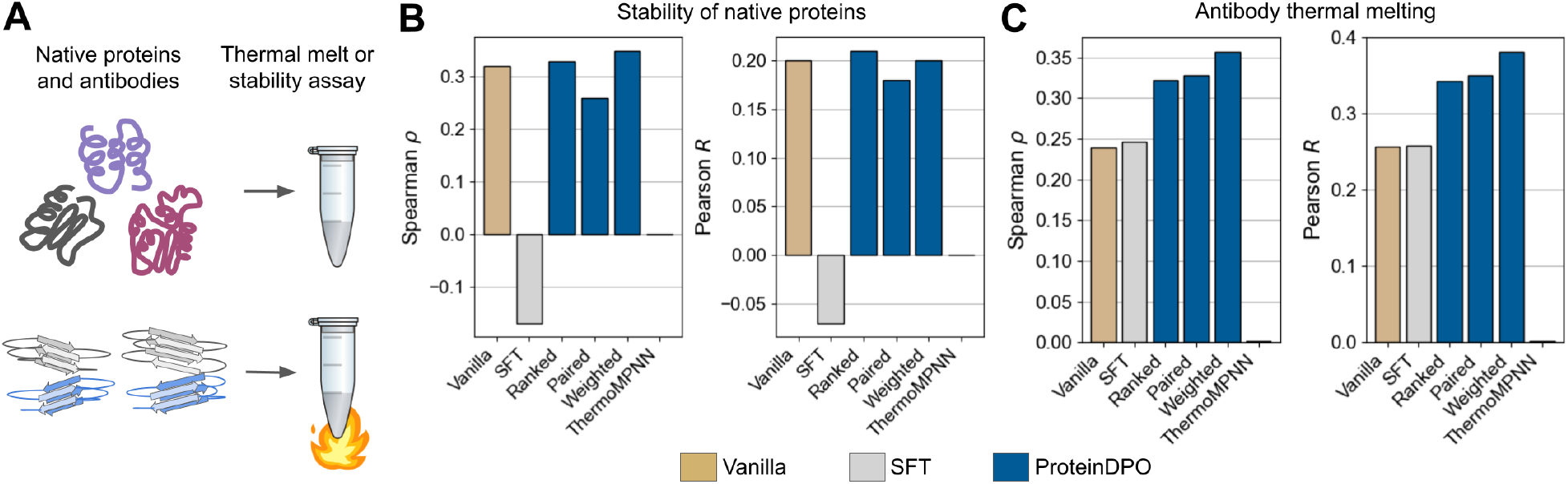
ProteinDPO generalizes to absolute stability prediction of diverse proteins. **(A)** We benchmarked models using datasets of absolute stability (ΔG) of natural proteins and thermal unfolding measurements (determination of the temperature which unfolds half the concentration of a protein) of antibodies. **(B)** Correlation of model likelihoods (Vanilla, SFT, ProteinDPO) or predicted values (ThermoMPNN) with experimentally measured stability of 26 native proteins from Maxwell et al. [31] **(C)** Correlation of model likelihoods with experimentally measured thermal melting points of clinical and naturally derived monoclonal antibodies (mAbs) curated from Shehata et al. [46] and Jain et al. [21].

Despite only training on relative stability data, ProteinDPO retains (**Figure 3B**) or improves (Pearson *R* and Spearman *ρ* increases of 0.08-0.12) on (**Figure 3C**) the ability of vanilla ESM-IF1 in ranking the absolute differences in free energy (ΔG) or thermal melting points (*T*_mid_) across natural proteins and antibodies [31] (**Figure 3A**). Meanwhile, the SFT model seems to have once again overfit to the training regime, completely losing the ability of vanilla ESM-IF1 to rank the absolute stability of natural proteins, and only performing similarly for the more homogeneous dataset of antibody thermostability. Notably, other supervised models trained on Megascale to predict ΔΔG (such as ThermoMPNN) cannot evaluate proteins with multiple chains (such as most antibodies) and absolute stability metrics of native sequences will, correctly, always have a predicted ΔΔG of zero.

### 2.4. Zero-shot binding affinity prediction

Motivated by previous work in which ESM-IF1 guided antibody affinity maturation by ranking mutant likelihoods conditioned on the respective antibody-antigen complex backbone [45], we were interested if our method, optimized for *stability*, will generalize to *binding affinity* given they are both determined by similar entropic and enthalpic interactions. To this end, we evaluate on two datasets, SKEMPIv2 [22] and AB-Bind [49], which consist of experimentally measured binding affinities for thousands of sequence variants across hundreds of protein-protein complexes (**Figure 4A**), with AB-Bind containing exclusively antibody-antigen complexes. Due to SKEMPIv2 containing very few variants per complex (as low as one) we report the correlation of all model likelihoods with all affinity measurements, while for AB-Bind we report correlations averaged across each complex. Surprisingly, despite only training on the ΔΔGs of small protein monomers, ProteinDPO has modestly increased scoring of the binding affinity of large protein-protein complexes (Pearson *R* increase of 0.02-0.04, Spearman *ρ* increase of 0.02-0.04, and AUROC increase of 0.04-0.05) (**Figure 4B**) and larger performance gains when scoring antibody-antigen complexes (Pearson *R* increase of 0.04-0.08, Spearman *ρ* increase of 0.03-0.10, AUROC increase of 0-0.02) (**Figure 4C**). Similar to previous results (**Figures 2,3**), the SFT model fails to generalize to binding-related tasks and, in some cases, actually results in worse-than-random performance (**Figure 3B**), consistent with SFT leading to overfitting.

**Figure 4.**
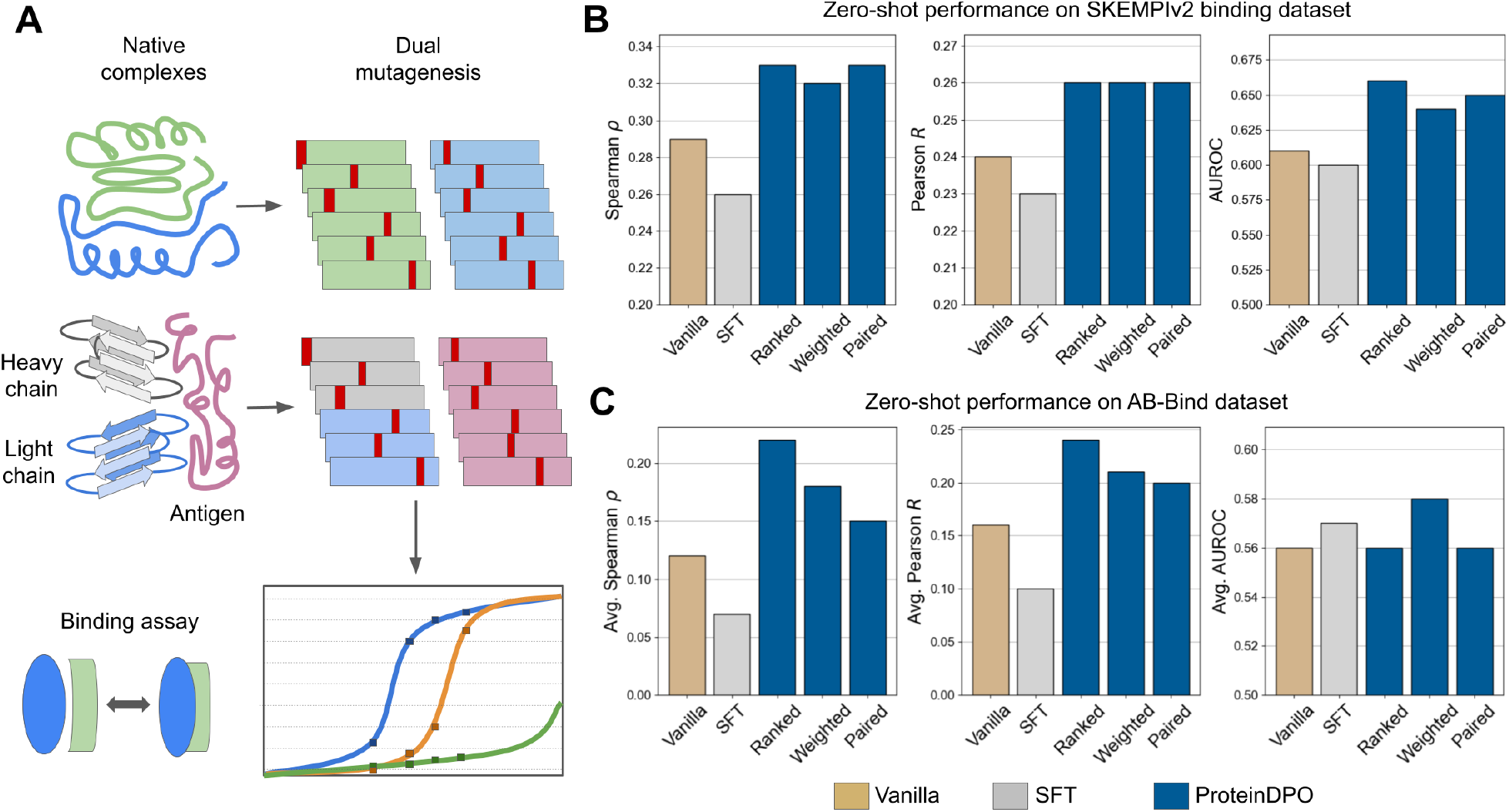
ProteinDPO generalizes to binding prediction. **(A)** We benchmarked models using binding affinity mutagenesis studies, which mutate one or multiple proteins that form a bound complex and subsequently measure the resulting binding affinity of the complex. **(B)** Zero-shot correlation and AUROC of model likelihoods with protein-protein binding affinity from the SKEMPIv2 dataset [22]. **(C)** Zero-shot correlation and AUROC of complex-conditioned model likelihoods with ΔΔG of antibody-antigen binding from the AB-Bind dataset [49], averaged across each complex. In **(B,C)**, the AUROC measures a model’s ability to classify mutants that increase or decrease affinity relative to the original complex.

### 2.5. Generation of stabilized proteins

Beyond merely a supervised scoring function, ProteinDPO is also a generative model. We were therefore interested in testing if ProteinDPO would be useful for stabilizing sequences for given protein backbones, and whether it could do so more favorably than the original ESM-IF1 model. This is a complex task, as combining favorable single-mutations to design a sequence that is more stable than the native protein is not trivial due to the non-linear and higher order interactions among protein residues [16].

For our generation experiments, we selected the “Paired” model due to its consistent performance across all scoring experiments. Here, we generate protein sequences for three diverse, native backbones (**Figure 5A**): staphylococcal nuclease [19] (PDB: 1STN), PGK [10] (PDB: 1PHP), and the membrane protein claudin-15 [53] (PDB: 4P79). We computed the distribution of the average Rosetta force-field energy (REU) across replicate runs for 500 generations for each model and native backbone, as well as the energy of each native sequence, all derived by modeling the mutated sequence onto the native backbone via the Rosetta *cart_ddg* protocol [13] (**Figure 5B**).

**Figure 5.**
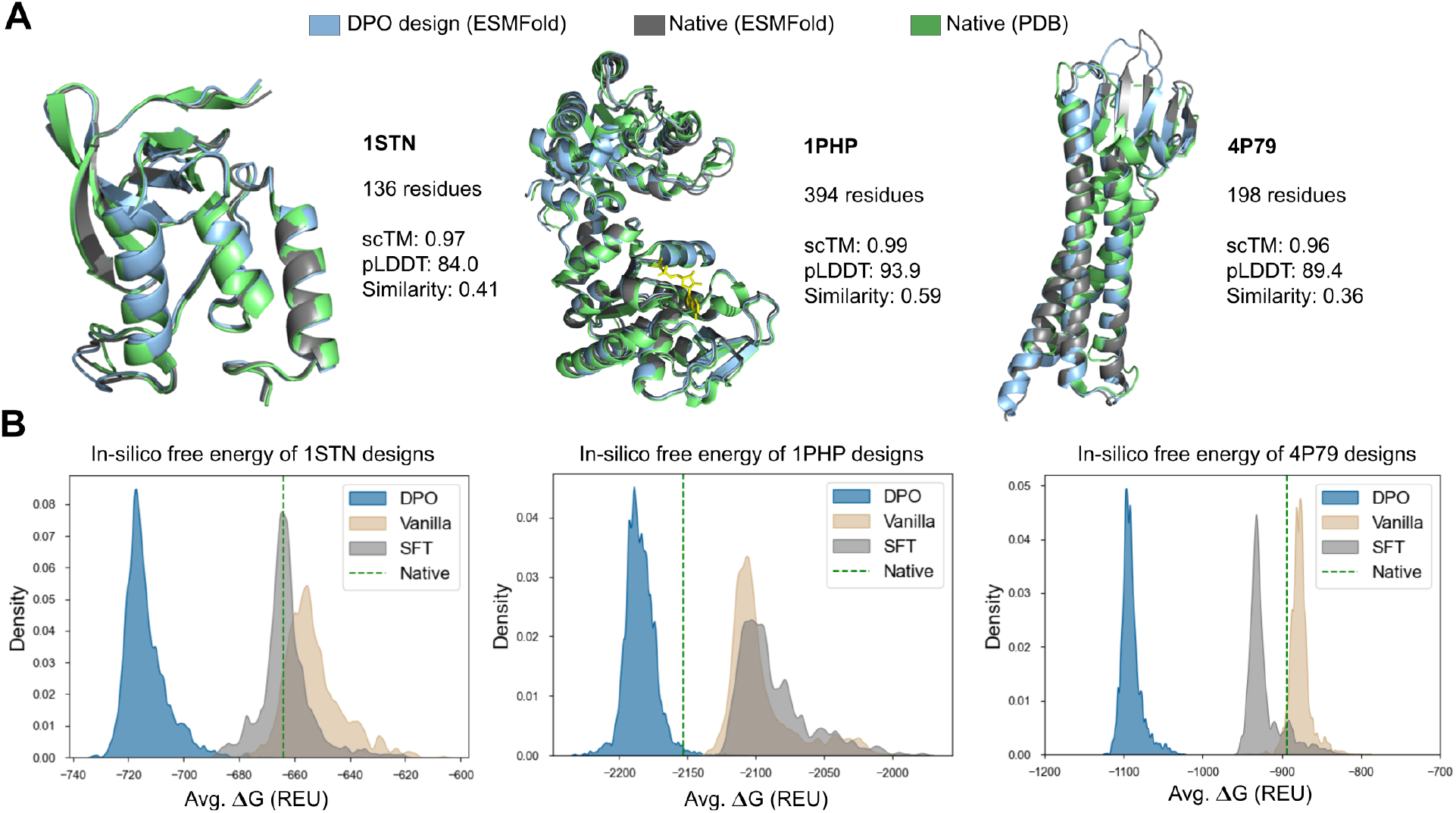
ProteinDPO generates stabilized proteins. **(A)** We evaluate the self-consistency and confidence of Paired model ProteinDPO designs for PDB structures 1STN, 1PHP and 4P79, comparing to experimental [5] and ESMFold predicted[28] structures. We show the experimental structure of the native sequence in green, the predicted structure of the native sequence in gray, and the predicted structure of the ProteinDPO design in blue. Self-consistency TM (scTM) is the structural similarity score (TM) of the native sequence predicted structure ProteinDPO sequence predicted structure; pLDDT is the average ESMFold [28] confidence (predicted error) for the structure of the generated sequence. Similarity is the percent of shared residue identity between the generation and native sequence. **(B)** Average *in-silico* free energy of protein folding (ΔG) for sequences from ProteinDPO, vanilla ESM-IF1, and SFT ESM-IF1 in Rosetta Energy Units (REU) compared to the native sequence derived from Rosetta *cart_ddg* protocol [13] with five replicate runs for each sequence. 500 samples were taken from each model across five sampling temperatures within [0.01, 1] with no additional filtering. Lower energy corresponds to higher stability.

Despite only training on the stabilities of small proteins (40 to 75 residues), sequences generated by ProteinDPO for much larger backbones (136 to 394 residues) are predicted to be substantially more stable (lower energy) than vanilla ESM-IF1, the SFT model, and the native sequence. Surprisingly, despite having significantly lower similarity to the native sequence than SFT and vanilla ESM-IF1, sequences generated by ProteinDPO have confident ESMFold-predicted structures (pLDDT > 80) (**Figure 5A**) that closely match the native structure (**Tables S1,S2,S3**). This suggests the stability optimized ProteinDPO has not lost the rules of structure recapitulation learned from the unsupervised pre-training of ESM-IF1.

## 3. Discussion

Here, we have shown that DPO is an effective way to imbue an unsupervised structure-conditioned language model with biophysical information. The resulting model, ProteinDPO, outperforms the original pretrained model and a model further specialized via supervised finetuning in its ability to score stability and to generate stable protein sequences. Strikingly, we see that the information learned from training ProteinDPO only on the relative changes in stability of small protein monomers generalizes to larger, unseen folds. This includes scoring the binding affinity of large protein-protein complexes, thermal melting points of multi-chain antibodies, and single-step stabilization of proteins an order of magnitude larger than the size of training data examples. In contrast, SFT, which does improve scoring within the training data, achieves poor generalization to these tasks, suggesting that SFT merely overfits to the data in the training distribution. Moreover, ProteinDPO’s generated sequences are still predicted to fold into the native structure, indicating that the model has not “forgotten” the information learned during unsupervised pretraining. In total, our results suggest that ProteinDPO has learned generalizable rules from its experimental biophysical alignment dataset.

While the model is competitive with supervised, task-specific models such as ThermoMPNN [11], it has not clearly surpassed all supervised models in scoring [8]. However, to our knowledge, ProteinDPO is the highest performing generative model at stability prediction, which enables stability-informed sequence generation for natural or de-novo protein folds with a single autoregressive sampling step. ProteinDPO also generalizes to a greater variety of tasks than existing supervised models, being able to predict ΔΔG, ΔG, binding affinity, and thermal melting over diverse wild-type proteins, multi-chain antibodies, and protein complexes. This is in contrast to models that are limited to scoring single-mutant ΔΔGs of monomeric structures, including ThermoMPNN [11].

While ProteinDPO incorporates stability information into a structure-conditioned language model, this approach generalizes to any labeled dataset of protein variants, to any generative model, and to biological data modalities beyond proteins. We find that alignment based on one specific property (such as the stability of small monomers) improves function prediction for other properties (such as binding affinity of large protein-protein complexes), most likely because both properties share common underlying physical principles. Future directions of this work could explore whether alignment based on other properties could likewise enable generalizable information transfer. We make ProteinDPO open-source and publicly available at https://github.com/evo-design/protein-dpo.

## 4. Data and code availability

We used the Megascale [55] dataset for training, either with our own training splits, those from Diekhaus et al. [11], or those from Cuturello et al. [8] depending on subsequent evaluation.

We used the following datasets for evaluation:

- Homologue-free FireProt [51, 11]
- S669 [36]
- Antibody thermal stability [46, 21]
- Native protein stabilites [31, 7]
- SKEMPIv2 [22]
- AB-Bind [49]

Additional details on these datasets are provided in **Methods**.

Code for ProteinDPO is available at https://github.com/evo-design/protein-dpo.

Weights for the model trained with the paired DPO objective are available at https://doi.org/10.5281/zenodo.11218181.

## 5. Methods

### 5.1. Generative model alignment

Generative AI applications are now routinely finetuned with feedback-based reinforcement learning (RL) during post-training, which has been crucial for the success of applications in the natural language domain [23, 15]. Here, we provide a brief overview of the classical model training pipeline, which consists of three stages: supervised finetuning, feedback collection, and RL finetuning [50, 35, 3].

#### Supervised finetuning (SFT)

Here, we define supervised finetuning as continuing the original pretraining objective on high-quality samples from a downstream labeled dataset. In this work for example, for supervised finetuning on the “Megascale” dataset, high-quality samples are sequences which achieve superior stability to the native sequence. During the SFT stage we use a dataset of input-output pairs, 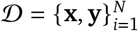 and train a model *p*_*θ*_ with next-token maximum likelihood loss on high-quality pairs, where *y*_*i*_ is the *i*^th^ token of the output **y**:

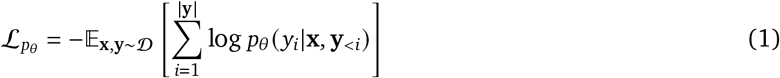

#### Feedback stage

During the feedback stage we use the base model or the SFT model *p*_SFT_ and a set of inputs to sample a number *K* > 0 of candidate completions **y**^(*i*)^, …, **y**^(*K*)^ per prompt **x** from a dataset of preferences 𝒟_pref_. Feedback can be provided in several ways:

1. Prior RLHF work has used ranking type of comparisons in which the candidate outputs are ranked 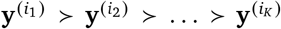, with pairwise rankings being a popular choice. In order to use this data from the reinforcement learning stage, we need to extract an evaluation score from it. We usually assume a parameterization of the preference distribution, such as the Plackett-Luce [38, 30] choice model (or the Bradley-Terry [6] for pairwise data). In the most general case, given the permutation 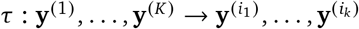, we have

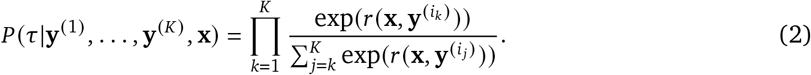

We can then formulate this as a maximum likelihood loss over the reward model

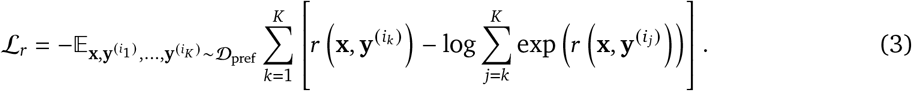
2. In the case of pairwise rankings **y**^*w*^ ≻ **y**^*l*^, this just reduces to the standard binary logistic regression loss,

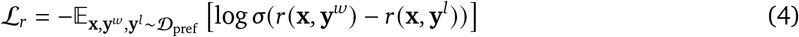

where *σ*(·) is the sigmoid function. Notice that if we are provided with a complete ranking over *K* candidates then we can construct 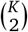 pairwise rankings, which was used in prior works [35]. These can still be easily optimized together since we still require a single forward pass through the reward model.
3. Alternatively a numerical score, *r*_*i*_ (such as used in a Likert scale) could be provided per candidate input-output pair (**x, y**^(*i*)^).

#### Reinforcement learning stage

During the reinforcement learning stage we use a reference model *p*_ref_, sometimes initialized as *p*_SFT_ from the SFT stage, and optimize the following RL problem:

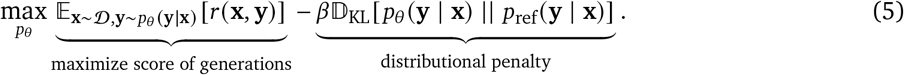

We want to optimize a model *p*_*θ*_ to maximize the score, while also maintaining a distribution close to the reference model. The second terms aims to keep the model close to the distribution, which was used to collect the feedback data and prevent out-of-distribution issues [27].

### 5.2. Optimization algorithms

Two main algorithms have emerged for the optimization of the objective in Eq. 5: (1) classical RL algorithms, such as PPO [44], which use on-policy rollouts from the model and (2) direct alignment algorithms, such as DPO [39], which train fully-offline using only feedback data. While successful in large scale-systems, such as GPT-4, the PPO pipeline requires a very accurate score model that can be queried in a continuous manner. Prior work has directly trained a separate score network *r*_*θ*_(**x, y**) using the feedback data with the objective in Eq. 4 and directly applied PPO to the problem in Eq. 5 using the trained score model. However, such a pipeline can be quite complex, unstable, and computationally expensive [59, 18]. DPO [39] is an alternative to this algorithmic pipeline, which we present below.

### 5.3. Direct Preference Optimization

The DPO algorithm is based on the observation that the objective in Eq. 5 has a closed form solution of the type

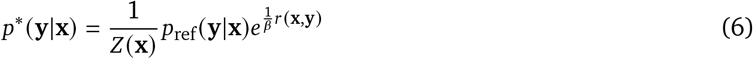

where

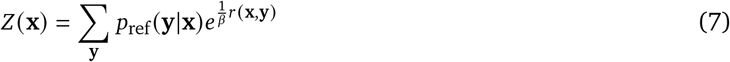

is an intractable partition function. The key insight of the DPO algorithm is the model-reward equivalency, i.e.,

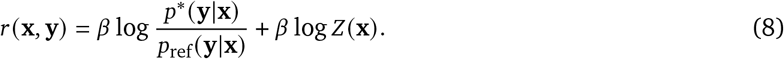

Now, if instead of training a score model *r*_*θ*_, we directly parameterize the generative model *p*_*θ*_, we can use the preference data to directly optimize *p*_*θ*_, essentially marginalizing the RL loop.

#### Ranked objective

If we substitute the above expression into the Plackett-Luce reward model in Eq. 3, we obtain the following objective:

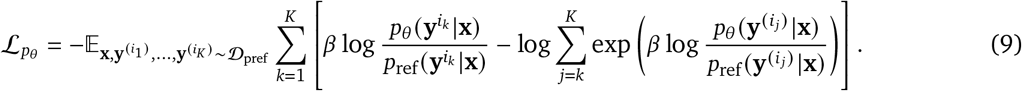

#### Paired objective

If we instead use the pairwise objective in Eq. 4, we obtain the canonical form of the DPO objective:

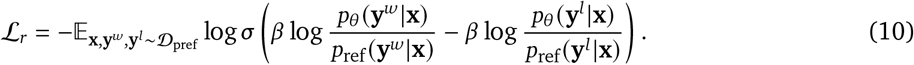

Here, due to the homogeneity of both the Plackett-Luce and Bradley-Terry model (i.e., comparing several options for the same context), the untractable partition term *Z*(**x**) cancels out and we are left with tractable, fully offline objectives.

#### 5.4. Weighted DPO algorithm

Prior methods, including DPO [39], have all focused on extracting a per-sample score *r* (**x, y**) from rankings. However, if we have some external evaluation score, i.e., data 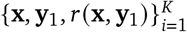, then this step is redundant.

Given numerical scores, we could train a score model, using a regression-based framework and then use the standard PPO pipeline, but this approach inherits all the shortcomings of the standard RLHF pipeline. We could use the numerical scores to induce a ranking over candidates **y**_*i*_ and the apply the Plackett-Luce or Bradley-Terry DPO objectives in Eq 9 and Eq 10 respectively, however that faces two drawbacks:

1. Plackett-Luce and Bradley-Terry both assume a particular parametric form of the ranking likelihood, i.e., the rankings are inherently probabilistic, which would induce additional noise in the optimization procedure if we generate rankings based on those choice models.
2. We can also generate rankings based on raw scores similarly to prior works [14]. However, note that any monotone transformation would still induce the same relative rankings. That is, there might be actual relative information in the score which could be lost in such a scenario.

Weighted regression-based methods [37, 32, 26] are a good fit for this problem, however, they have been sub-optimal in the large generative model setting and do not yield significant improvement [54]. Instead, we will aim to develop a DPO type of algorithm that can efficiently utilize the numerical scores of the candidate sequences. Given an **x** and candidate sequences 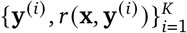, consider the parameterized reward

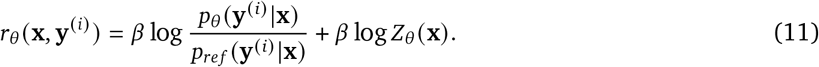

That is a parameterized distribution *p*_*θ* (**y**|**x**)_ induces a reward function *r*_*θ*_ (**x, y**), such that the model *p*_*θ*_ is the optimal solution for the corresponding RL problem in Eq. 5. Now, instead of aligning the generative model *p*_*θ*_ with some sampled rankings or preferences, we will instead minimize a distance function of the type *D*(*r*(**x, y**), *r*_*θ*_ (**x, y**)); that is, we will match the true score samples with the implicit ones. A remaining issue is that we still have the constraining partition function *Z*_*θ*_ (**x**) (since it depends on *p*_*θ*_), which we would like to eliminate. When we consider

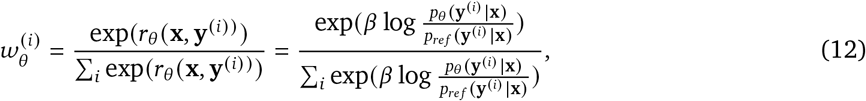

notice that the partition function *Z*_*θ*_ (**x**) was eliminated. In the same manner we have that

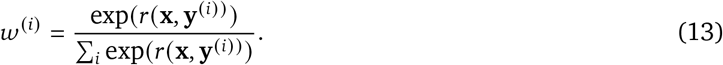

Notice now that **w** = (***w***^(1)^, …, ***w***^(*K*)^) and 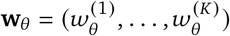 are both valid probability distributions. In fact, the probability distributions resemble the Boltzmann Distribution from statistical mechanics [34],

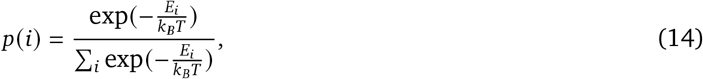

where *p*(*i*) is the probability of the system being in a state *i* with energy *E*_*i*_, and *k*_*B*_*T* is the scaled absolute temperature of the system. Substituting *r*(**x, y**^(*i*)^) for the negative energy of the state −*E*_*i*_, and considering *k*_*B*_*T* a parameter which allows us to scale the numerical scores *r* (**x, y**^(*i*)^), the weighted DPO objective is simply minimization of the KL-divergence 𝔻_*KL*_[**w**|**w**_*θ*_] between the Boltzmann-like distributions parameterized by the generative model *p*_*θ*_ and numerical scores from the dataset 𝒟. With some simple algebra, this yields:

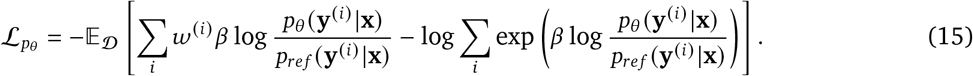

In the case where we have only two candidate sequences **y**^(1)^ and **y**^(2)^, this can be simplified to

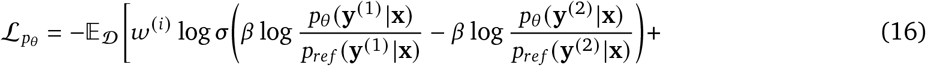

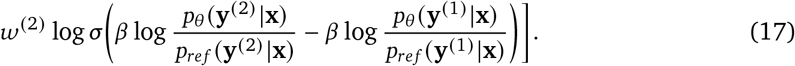

That is, this is essentially a weighted version of the DPO algorithm. One interpretation is that this objective is the expected DPO objective under *p*(**y**^(1)^ ≻ **y**^(2)^ |**x**) = ***w***^(1)^ and *p*(**y**^(2)^ ≻ **y**^(1)^ |**x**) = ***w***^(2)^. These objectives are theoretically optimal in the offline regime as they are both mode-seeking and use a “negative gradient” formulation [54].

### 5.5. Datasets

#### Megascale

The Megascale dataset served as the experimentally measured stability dataset from which we aligned the pretrained ESM-IF1. After filtering out data marked as unreliable for machine learning by Megascale authors, the dataset contained approximately 660k variants for 405 protein domains. The variants consisted of insertions, deletions, single and double substitutions. Similar to prior methods [11], insertions and deletions were not considered as DPO training was designed to compare stabilities across the same structure, however due to the robust nature of the model we were able to train and model with the double-mutations in the dataset. We additionally removed duplicate sequences. FoldSeek [56] clustering with the “easy-cluster” algorithm was performed on all PDBs remaining in the Megascale dataset with an alignment threshold of 50%. From this, clusters were randomly assigned to respective train (90%), validation (5%), and test (5%) datasets to ensure generalizability of the model across different structures. Because the model is trained with sets of sequences for a common structure, either the ΔG (absolute) or ΔΔG (relative) of sequences were used for training. While these should be identical in theory after accounting for native ΔG, due to noise in the data the two metrics are different. We found negligible difference in loss and validation accuracy training with either metric, so all models were trained on ΔG as more data points had ΔG measurements. For supervised finetuning, all variants with ΔΔG < 0 were selected and the same splits were used.

#### SKEMPIv2

SKEMPIv2 is database of binding free energy changes upon single point mutations within a variety of protein complex interfaces [22]. After removal of PDBs with non-canonical amino acids, the final dataset contained 6,487 variants. To score on this dataset, we followed instructions and code from the ESM-IF1 Github [17], averaging the log-likelihood of simultaneous mutations given the entire complex structure and sequence as input. However we took the additional step of normalizing with the averaged wild-type likelihood of said mutated residues. Because certain complexes were only represented by very few variants, sometimes with a different PDB for each, we did not evaluate the average correlation within each complex, but report an “all vs. all” comparison.

**Figure S1.**
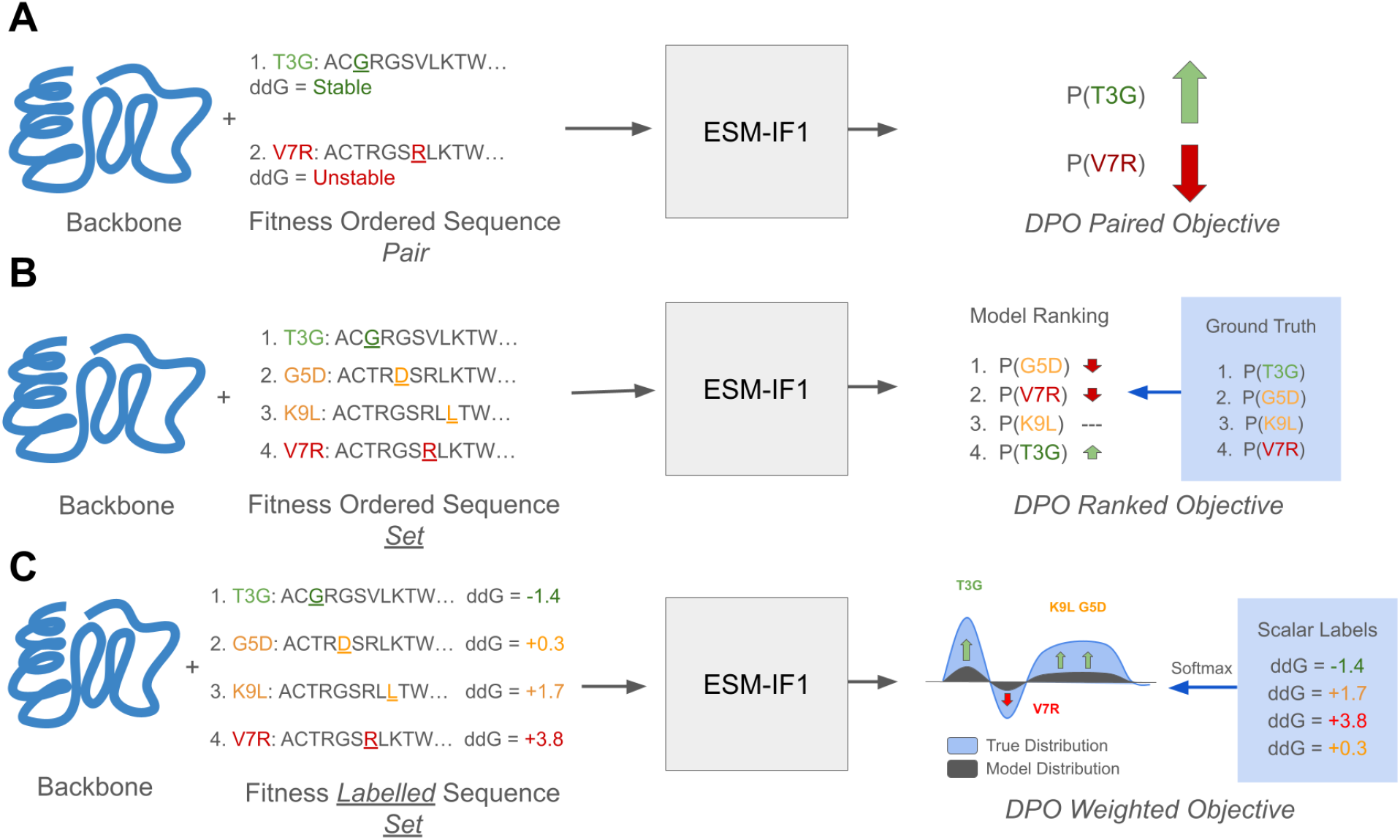
Input Construction and Objectives across DPO Reward Models. **(A)** The paired reward model is the standard algorithm which satisfies the DPO objective under the Bradley-Terry model for reward. For a given backbone prompt, the loss function takes in an ordered pair of sequences responses with “good” and “bad” stability, and will train the model to prefer the “good” sequence of the “bad” sequence for the given backbone. **(B)** The ranked reward model satisfies the DPO objective under the Plackett-Luce model for reward. For a given backbone prompt, the loss function takes in an ordered set of sequence responses, and will train the model to prefer sequences in the same respective ranking. **(C)** The weighted reward model will, given a backbone prompt, accept a set of sequences labeled with fitness scores, and will train the model’s distribution of likelihood over the sequences to match the relative distribution of reward of the sequences derived from the softmax of their scalar labels.

#### AB-Bind

AB-Bind is an antibody binding mutational database curated to train and evaluate computational affinity predictions from Sirin et al. [49]. The database includes 1,101 mutants across 32 complexes curated from public data, each mutant having experimentally determined changes in binding free energy (ΔΔG). Duplicated variants, variants with missing mutated residues in the PDB, and those without an experimentally determined PDB were removed resulting in a final dataset size of 1,011 variants. We evaluated scoring on this dataset the same as with SKEMPIv2 [22], however we reported the average correlations across each antibody-antigen complex as there were an ample number of variants per complex.

#### Antibody thermal unfolding

The antibody thermostability dataset is derived from Widatalla et al. [58] which compiled scores and IgFold [42] predicted structures from the original Jain et al. [21] and Shehata et al. [46]datasets, consisting of thermostability measurements for monoclonal antibododies (mAbs). These two studies collected several biophysical measurements for 500 human B-Cell and LLPC derived mAbs [46], and mAbs that have been in clinical trials [21]. Among these includes 483 measurements of the temperature of thermal unfolding (T-mid). While the entire fragment antigen binding (Fab) domain was measured in this experiment, only the structure and sequence of the variable domains (Fv) were taken as input to the models. We used no normalization for scoring on this dataset as we considered every sequence as a “native” protein.

#### Native protein stabilities

The dataset of native protein stabilities from Maxwell et. al [31] is an experimentally measured dataset of the free energies (ΔGs) of unfolding for two-state proteins whose kinetics and thermodynamics have been studied biophysically, and that range in size from 61–151 residues. We used the curated version of this dataset from Cagiada et al. [7]. containing 26 proteins. Following Cagiada et al. [7], we used the sum of the log-likelihoods as opposed to the average due to the size dependence of absolute protein free energy.

#### FireProt

The FireProt database is a curated dataset of changes in free energy (ΔΔG) for 3,438 single mutations for 100 unique proteins [51]. To alleviate data-leakage from the Megascale training data, we used the Megascale “Homologue-Free” FireProt dataset from Diekhaus et al. [11] which has all homologous sequences to the Megascale dataset removed, resulting in 2,578 variants. For this dataset, likely due to the presence of only single mutations, we found the best performance taking the log-likelihood of only the mutated residue, and normalizing by subtracting the wild-type residue likelihood.

#### S669

The S669 dataset contains experimentally measured ΔΔG values of 669 single mutations of 94 proteins [36]. Due to the presence of non-canonical amino acids, 4 variants were removed from this dataset for evaluation of vanilla ESM-IF1 and ProteinDPO. Similar to FireProt [51], we found the best performance taking the log likelihood of the mutated residue normalized by subtracting the wild-type residue likelihood. Results for all other methods on this dataset derived from Cuturello et al. and Diekhaus et al. [8, 11].

### 5.6. Model training

#### Input construction

Multiple methods for positive-negative pair selection from the scalar-labeled data were evaluated, including randomly selecting pairs then ordering them by scalar value, prioritizing pairs with small differences, and prioritizing pairs with large differences. To have a tunable parameter that controls the distance between pairs for the two latter approaches, we implemented a “gap level” heuristic, which is a parameter within [0,1]. A gap level of 1 corresponds to placing the largest gap possible in sequence ranking between pairs, while a gap level of 0 corresponds to the smallest possible gap. For any gap level choice, we ensure the gaps between sequences in terms of ranking is uniform across all pairs, and that all sequences are used, except for a single excluded sequence in the case there is an odd number of sequences for a native protein. While most choices of gap level had comparable performance, we found a gap level of 0.5 performed the best on validation data. Sets of sequences were randomly chosen for models trained with the ranked objective, and models trained with the weighted objective and set size (*K*) greater than two. For models trained with the weighted objective we experimented with various values temperature values (*T*) for scaling the scalar labels, finding 0.1 to be the most favorable (**Table S5**).

#### Training details

Noise of 0.1 Angstroms was applied to all PDB structures in line with the training of ESM-IF1 [17]. For each set (Ranked and Weighted models) or pair (Paired model) of sequence responses, the same native backbone was used as structural prompt, PDBs of which were derived from Tsuboyama et al. [55]. Hyperparameter tuning was performed with varying values for *β*, gap level (Paired and Weighted Model), temperature *T* (Weighted Model), learning rate, *K* (Weighted and Ranked Model) and batch size. For computational efficiency, reference model likelihoods were pre-computed before training. Final hyperparameters for the Ranked, Paired and Weighted models are provided in **Table S5**. Models were each trained until validation loss convergence with the AdamW [29] optimizer on a single NVIDIA H100 GPU with 80GB of memory.

## 6. Acknowledgements

The authors would like to thank Chiho Im, Richard Shuai, Brian Kang, and Garyk Brixi for helpful discussions and Jeremy Sullivan for cluster support. T.W. is supported by the NSF Graduate Research Fellowship and the Stanford Graduate Fellowship. B.L.H. acknowledges funding support from Arc Institute.

## 7. Author contributions

T.W. and B.L.H, conceived of the research. T.W. performed dataset curation, model training, generations and evaluation. R.R. derived the DPO algorithms. B.L.H. supervised the research. T.W. wrote the initial version of the manuscript. All authors wrote the final manuscript.

## 9. Supplemental

**Table S1.**
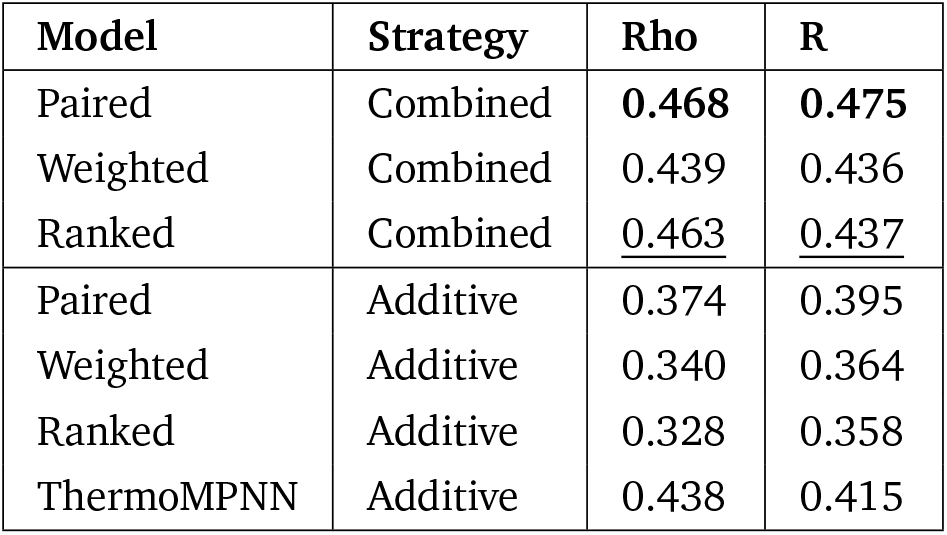
Additive and whole-sequence scoring on Megascale holdout double mutants.

**Table S2.** Self-consistency of 4P79 generations.

| Temp | ProteinDPO |  |  | Vanilla |  |  | SFT |  |  |
| --- | --- | --- | --- | --- | --- | --- | --- | --- | --- |
|  | scTM | pLDDT | %Sim | scTM | pLDDT | %Sim | scTM | pLDDT | %Sim |
| 0.01 | 0.96 | 90.2 | 37.5 | 0.95 | 88.8 | 45.5 | 0.94 | 80.5 | 41.8 |
| 0.05 | 0.96 | 90.6 | 37.9 | 0.95 | 88.8 | 45.2 | 0.93 | 81.1 | 41.4 |
| 0.1 | 0.96 | 90.6 | 37.7 | 0.95 | 88.0 | 45.2 | 0.93 | 82.1 | 41.5 |
| 0.5 | 0.96 | 89.3 | 35.5 | 0.95 | 85.6 | 45.7 | 0.94 | 82.0 | 39.4 |
| 1.0 | 0.95 | 86.2 | 33.4 | 0.94 | 82.4 | 41.5 | 0.94 | 82.5 | 34.7 |

**Table S3.**
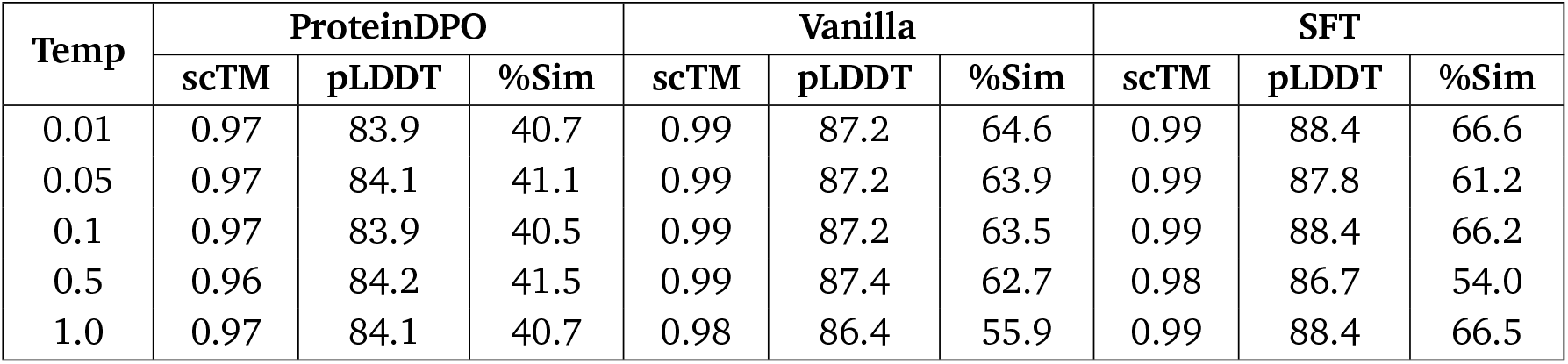
Self-consistency of 1STN generations.

**Table S4.** Self-consistency of 1PHP generations.

| Temp | ProteinDPO |  |  | Vanilla |  |  | SFT |  |  |
| --- | --- | --- | --- | --- | --- | --- | --- | --- | --- |
|  | scTM | pLDDT | %Sim | scTM | pLDDT | %Sim | scTM | pLDDT | %Sim |
| 0.01 | 0.99 | 93.4 | 60.9 | 0.99+ | 90.1 | 79.8 | 0.99+ | 91.4 | 75.2 |
| 0.05 | 0.99 | 93.8 | 58.6 | 0.99+ | 91.3 | 77.4 | 0.99+ | 92.2 | 73.7 |
| 0.1 | 0.99 | 93.5 | 61.2 | 0.99+ | 89.9 | 80.0 | 0.99+ | 91.0 | 75.0 |
| 0.5 | 0.99 | 95.0 | 55.1 | 0.99+ | 93.1 | 69.9 | 0.99+ | 92.6 | 67.5 |
| 1.0 | 0.99 | 93.8 | 60.4 | 0.99+ | 90.1 | 79.4 | 0.99+ | 91.4 | 75.6 |

**Table S5.** Details on ProteinDPO model hyperparameters.

| Model | Paired | Ranked | Weighted |
| --- | --- | --- | --- |
| Learning Rate | $1 \times 10^{-7}$ | $1 \times 10^{-7}$ | $1 \times 10^{-7}$ |
| Batch size | 32 | 8 | 32 |
| $K$ | 2 | 3 | 2 |
| Gap level | 0.5 | N/A | 0.5 |
| $T$ | N/A | N/A | 0.1 |
| $\beta$ | 0.1 | 0.1 | 0.1 |
| Max epochs | 30 | 30 | 30 |
| Adam $\beta_1, \beta_2$ | 0.9, 0.98 | 0.9, 0.98 | 0.9, 0.98 |
| Adam $\epsilon$ | $1 \times 10^{-8}$ | $1 \times 10^{-8}$ | $1 \times 10^{-8}$ |
| Adam decay | 0.1 | 0.1 | 0.1 |

## References

[1] Anishchenko, S. J. Pellock, T. M. Chidyausiku, T. A. Ramelot, S. Ovchinnikov, J. Hao, K. Bafna, C. Norn, A. Kang, A. K. Bera, and et al. De novo protein design by deep network hallucination. Nature, 600(7889):547–552, 2021.

[2] M. Baek, F. Dimaio, I. Anishchenko, J. Dauparas, S. Ovchinnikov, G. R. Lee, J. Wang, Q. Cong, L. N. Kinch, R. D. Schaeffer, and et al. Accurate prediction of protein structures and interactions using a three-track neural network. Science, 373(6557):871–876, 2021.

[3] Y. Bai, A. Jones, K. Ndousse, A. Askell, A. Chen, N. DasSarma, D. Drain, S. Fort, D. Ganguli, T. Henighan, N. Joseph, S. Kadavath, J. Kernion, T. Conerly, S. El-Showk, N. Elhage, Z. Hatfield-Dodds, D. Hernandez, T. Hume, S. Johnston, S. Kravec, L. Lovitt, N. Nanda, C. Olsson, D. Amodei, T. Brown, J. Clark, S. McCandlish, C. Olah, B. Mann, and J. Kaplan. Training a helpful and harmless assistant with reinforcement learning from human feedback, 2022.

[4] M. Bailly, C. Mieczkowski, V. Juan, E. Metwally, D. Tomazela, J. Baker, M. Uchida, E. Kofman, F. Raoufi, S. Motlagh, and et al. Predicting antibody developability profiles through early stage discovery screening. mAbs, 12(1):1743053, 2020.

[5] H. M. Berman, J. Westbrook, Z. Feng, G. Gilliland, T. N. Bhat, H. Weissig, I. N. Shindyalov, and P. E. Bourne. The Protein Data Bank. Nucleic Acids Research, 28(1):235–242, 01 2000.

[6] R. A. Bradley and M. E. Terry. Rank analysis of incomplete block designs: I. the method of paired comparisons. Biometrika, 39(3/4):324–345, 1952.

[7] M. Cagiada, S. Ovchinnikov, and K. Lindorff-Larsen. Predicting absolute protein folding stability using generative models. bioRxiv, 2024.

[8] F. Cuturello, M. Celoria, A. Ansuini, and A. Cazzaniga. Enhancing predictions of protein stability changes induced by single mutations using msa-based language models. bioRxiv, 2024.

[9] J. Dauparas, I. Anishchenko, N. Bennett, H. Bai, R. J. Ragotte, L. F. Milles, B. I. M. Wicky, A. Courbet, R. J. de Haas, N. Bethel, P. J. Y. Leung, T. F. Huddy, S. Pellock, D. Tischer, F. Chan, B. Koepnick, H. Nguyen, Kang, B. Sankaran, A. K. Bera, N. P. King, and D. Baker. Robust deep learning–based protein sequence design using proteinmpnn. Science, 378(6615):49–56, 2022.

[10] G. J. Davies, S. J. Gamblin, J. A. Littlechild, Z. Dauter, K. S. Wilson, and H. C. Watson. Structure of the ADP complex of the 3-phosphoglycerate kinase from Bacillus stearothermophilus at 1.65 Å. Acta Crystallographica Section D, 50(2):202–209, Mar 1994.

[11] H. Dieckhaus, M. Brocidiacono, N. Z. Randolph, and B. Kuhlman. Transfer learning to leverage larger datasets for improved prediction of protein stability changes. Proceedings of the National Academy of Sciences, 121(6):e2314853121, 2024.

[12] Doerr. Widening the protein crystallization bottleneck. Nature Methods, 3(12):961–961, 2006.

[13] Frenz, S. M. Lewis, I. King, F. Dimaio, H. Park, and Y. Song. Prediction of protein mutational free energy: Benchmark and sampling improvements increase classification accuracy. Frontiers in Bioengineering and Biotechnology, 8, 2020.

[14] L. Gao, J. Schulman, and J. Hilton. Scaling laws for reward model overoptimization. International Conference on machine Learning, 2023.

[15] M. GenAI. Introducing meta llama 3: The most capable openly available llm to date, 2024.

[16] M. Gromiha and S. Selvaraj. Inter-residue interactions in protein folding and stability. Progress in Biophysics and Molecular Biology, 86(2):235–277, 2004.

[17] C. Hsu, R. Verkuil, J. Liu, Z. Lin, B. Hie, T. Sercu, A. Lerer, and A. Rives. Learning inverse folding from millions of predicted structures. In K. Chaudhuri, S. Jegelka, L. Song, C. Szepesvari, G. Niu, and S. Sabato, editors, Proceedings of the 39th International Conference on Machine Learning, volume 162 of Proceedings of Machine Learning Research, pages 8946–8970. PMLR, 17–23 Jul 2022.

[18] S. Huang, R. F. J. Dossa, A. Raffin, A. Kanervisto, and W. Wang. The 37 implementation details of proximal policy optimization. In ICLR Blog Track, 2022. https://iclr-blog-track.github.io/2022/03/25/ppo-implementation-details/.

[19] T. R. Hynes and R. O. Fox. The crystal structure of staphylococcal nuclease refined at 1.7 Å resolution. Proteins: Structure, Function, and Bioinformatics, 10(2):92–105, 1991.

[20] J. Ingraham, V. Garg, R. Barzilay, and T. Jaakkola. Generative models for graph-based protein design. In H. Wallach, H. Larochelle, A. Beygelzimer, F. d’Alché-Buc, E. Fox, and R. Garnett, editors, Advances in Neural Information Processing Systems, volume 32. Curran Associates, Inc., 2019.

[21] T. Jain, T. Sun, S. Durand, A. Hall, N. R. Houston, J. H. Nett, B. Sharkey, B. Bobrowicz, I. Caffry, Y. Yu, Y. Cao, H. Lynaugh, M. Brown, H. Baruah, L. T. Gray, E. M. Krauland, Y. Xu, M. Vásquez, and K. D. Wittrup. Biophysical properties of the clinical-stage antibody landscape. Proceedings of the National Academy of Sciences, 114(5):944–949, 2017.

[22] J. Jankauskaitė, B. Jiménez-García, J. Dapkūnas, J. Fernández-Recio, and I. H. Moal. SKEMPI 2.0: an updated benchmark of changes in protein–protein binding energy, kinetics and thermodynamics upon mutation. Bioinformatics, 35(3):462–469, 07 2018.

[23] B. Z. John Schulman, C. Kim, J. Hilton, J. Menick, J. Weng, J. F. C. Uribe, L. Fedus, M. P. Luke Metz, R. G. Lopes, S. Zhao, A. Vijayvergiya, E. Sigler, A. Perelman, C. Voss, M. Heaton, J. Parish, D. Cummings, R. Nayak, V. Balcom, D. Schnurr, T. Kaftan, C. Hallacy, N. Turley, N. Deutsch, V. Goel, J. Ward, A. Konstantinidis, W. Zaremba, L. Ouyang, L. Bogdonoff, J. Gross, D. Medina, S. Yoo, T. Lee, R. Lowe, D. Mossing, J. Huizinga, R. Jiang, C. W. amd Diogo Almeida, S. Lin, M. Zhang, K. Xiao, K. Slama, S. Bills, A. Gray, J. Leike, J. Pachocki, P. Tillet, S. Jain, G. Brockman, N. Ryder, A. Paino, Q. Yuan, C. Winter, B. Wang, M. Bavarian, I. Babuschkin, S. Sidor, I. Kanitscheider, M. Pavlov, M. Plappert, N. Tezak, H. Jun, W. Zhuk, V. Pong, L. Kaiser, J. Tworek, A. Carr, L. Weng, S. Agarwal, K. Cobbe, V. Kosaraju, A. Power, S. Polu, J. Han, R. Puri, S. Jain, B. Chess, C. Gibson, O. Boiko, E. Parparita, A. Tootoonchian, K. Kosic, and C. Hesse. Introducing chatgpt, 2022.

[24] J. Jumper, R. Evans, A. Pritzel, T. Green, M. Figurnov, O. Ronneberger, K. Tunyasuvunakool, R. Bates, A. Žídek, A. Potapenko, and et al. Highly accurate protein structure prediction with alphafold. Nature, 596(7873):583–589, 2021.

[25] M. Knudsen and C. Wiuf. The cath database. Human Genomics, 4(3):207, 2010.

[26] Kostrikov, A. Nair, and S. Levine. Offline reinforcement learning with implicit q-learning, 2021.

[27] N. Lambert and R. Calandra. The alignment ceiling: Objective mismatch in reinforcement learning from human feedback, 2023.

[28] Z. Lin, H. Akin, R. Rao, B. Hie, Z. Zhu, W. Lu, N. Smetanin, R. Verkuil, O. Kabeli, Y. Shmueli, A. dos Santos Costa, M. Fazel-Zarandi, T. Sercu, S. Candido, and A. Rives. Evolutionary-scale prediction of atomic-level protein structure with a language model. Science, 379(6637):1123–1130, 2023.

[29] Loshchilov and F. Hutter. Decoupled weight decay regularization, 2019.

[30] R. D. Luce. Individual choice behavior: A theoretical analysis. Courier Corporation, 2012.

[31] L. Maxwell, D. Wildes, A. Zarrine-Afsar, M. A. De Los Rios, A. G. Brown, C. T. Friel, L. Hedberg, J.-C. Horng, D. Bona, E. J. Miller, A. Vallée-Bélisle, E. R. Main, F. Bemporad, L. Qiu, K. Teilum, N.-D. Vu, A. M. Edwards, I. Ruczinski, F. M. Poulsen, B. B. Kragelund, S. W. Michnick, F. Chiti, Y. Bai, S. J. Hagen, L. Serrano, M. Oliveberg, D. P. Raleigh, P. Wittung-Stafshede, S. E. Radford, S. E. Jackson, T. R. Sosnick, S. Marqusee, A. R. Davidson, and K. W. Plaxco. Protein folding: Defining a “standard” set of experimental conditions and a preliminary kinetic data set of two-state proteins. Protein Science, 14(3):602–616, 2005.

[32] Nair, A. Gupta, M. Dalal, and S. Levine. Awac: Accelerating online reinforcement learning with offline datasets, 2021.

[33] E. Nijkamp, J. A. Ruffolo, E. N. Weinstein, N. Naik, and A. Madani. Progen2: Exploring the boundaries of protein language models. Cell Systems, 14(11):968–978.e3, 2023.

[34] F. Noé, S. Olsson, J. Köhler, and H. Wu. Boltzmann generators: Sampling equilibrium states of many-body systems with deep learning. Science, 365(6457):eaaw1147, 2019.

[35] Ouyang, J. Wu, X. Jiang, D. Almeida, C. Wainwright, P. Mishkin, C. Zhang, S. Agarwal, K. Slama, A. Ray, J. Schulman, J. Hilton, F. Kelton, L. Miller, M. Simens, A. Askell, P. Welinder, P. F. Christiano, J. Leike, and R. Lowe. Training language models to follow instructions with human feedback. In S. Koyejo, S. Mohamed, A. Agarwal, D. Belgrave, K. Cho, and A. Oh, editors, Advances in Neural Information Processing Systems, volume 35, pages 27730–27744. Curran Associates, Inc., 2022.

[36] C. Pancotti, S. Benevenuta, G. Birolo, V. Alberini, V. Repetto, T. Sanavia, E. Capriotti, and P. Fariselli. Predicting protein stability changes upon single-point mutation: a thorough comparison of the available tools on a new dataset. Briefings in Bioinformatics, 23(2):bbab555, 01 2022.

[37] X. B. Peng, A. Kumar, G. Zhang, and S. Levine. Advantage-weighted regression: Simple and scalable off-policy reinforcement learning. arXiv preprint arXiv:1910.00177, 2019.

[38] R. L. Plackett. The analysis of permutations. Journal of the Royal Statistical Society. Series C (Applied Statistics), 24(2):193–202, 1975.

[39] R. Rafailov, A. Sharma, E. Mitchell, C. D. Manning, S. Ermon, and C. Finn. Direct preference optimization: Your language model is secretly a reward model. In Thirty-seventh Conference on Neural Information Processing Systems, 2023.

[40] G. J. Rocklin, T. M. Chidyausiku, I. Goreshnik, A. Ford, S. Houliston, A. Lemak, L. Carter, R. Ravichandran, V. K. Mulligan, A. Chevalier, C. H. Arrowsmith, and D. Baker. Global analysis of protein folding using massively parallel design, synthesis, and testing. Science, 357(6347):168–175, 2017.

[41] Z. A. Rollins, T. Widatalla, A. Waight, A. C. Cheng, and E. Metwally. Ablef: antibody language ensemble fusion for thermodynamically empowered property predictions. Bioinformatics, 40(5), 2024.

[42] A. Ruffolo, L.-S. Chu, S. P. Mahajan, and J. J. Gray. Fast, accurate antibody structure prediction from deep learning on massive set of natural antibodies. Nature Communications, 14(1), 2023.

[43] R. Schmirler, M. Heinzinger, and B. Rost. Fine-tuning protein language models boosts predictions across diverse tasks. bioRxiv, 2023.

[44] Schulman, F. Wolski, P. Dhariwal, A. Radford, and O. Klimov. Proximal policy optimization algorithms, 2017.

[45] V. R. Shanker, T. U. Bruun, B. L. Hie, and P. S. Kim. Inverse folding of protein complexes with a structure-informed language model enables unsupervised antibody evolution, 2023.

[46] Shehata, D. P. Maurer, A. Z. Wec, A. Lilov, E. Champney, T. Sun, K. Archambault, I. Burnina, H. Lynaugh, X. Zhi, and et al. Affinity maturation enhances antibody specificity but compromises conformational stability. Cell Reports, 28(13):3300–3308.e4, 2019.

[47] D. Shortle. One sequence plus one mutation equals two folds. Proceedings of the National Academy of Sciences, 106(50):21011–21012, 2009.

[48] R. W. Shuai, J. A. Ruffolo, and J. J. Gray. Iglm: Infilling language modeling for antibody sequence design. Cell Systems, 14(11):979–989.e4, 2023.

[49] S. Sirin, J. R. Apgar, E. M. Bennett, and A. E. Keating. Ab-bind: Antibody binding mutational database for computational affinity predictions. Protein Science, 25(2):393–409, 2016.

[50] Stiennon, L. Ouyang, J. Wu, D. M. Ziegler, R. Lowe, C. Voss, A. Radford, D. Amodei, and P. Christiano. Learning to summarize from human feedback, 2022.

[51] J. Stourac, J. Dubrava, M. Musil, J. Horackova, J. Damborsky, S. Mazurenko, and D. Bednar. FireProtDB: database of manually curated protein stability data. Nucleic Acids Research, 49(D1):D319–D324, 11 2020.

[52] H. Sumida, R. Núñez-Franco, I. Kalvet, S. J. Pellock, B. I. M. Wicky, L. F. Milles, J. Dauparas, J. Wang, Y. Kipnis, N. Jameson, A. Kang, J. De La Cruz, B. Sankaran, A. K. Bera, G. Jiménez-Osés, and D. Baker. Improving protein expression, stability, and function with proteinmpnn. Journal of the American Chemical Society, 146(3):2054–2061, 2024. PMID: 38194293.

[53] H. Suzuki, T. Nishizawa, K. Tani, Y. Yamazaki, A. Tamura, R. Ishitani, N. Dohmae, S. Tsukita, O. Nureki, and Y. Fujiyoshi. Crystal structure of a claudin provides insight into the architecture of tight junctions. Science, 344(6181):304–307, 2014.

[54] F. Tajwar, A. Singh, A. Sharma, R. Rafailov, J. Schneider, T. Xie, S. Ermon, C. Finn, and A. Kumar. Preference fine-tuning of llms should leverage suboptimal, on-policy data, 2024.

[55] K. Tsuboyama, J. Dauparas, J. Chen, E. Laine, Y. Mohseni Behbahani, J. J. Weinstein, N. M. Mangan, S. Ovchinnikov, and G. J. Rocklin. Mega-scale experimental analysis of protein folding stability in biology and design. Nature, 620(7973):434–444, 2023.

[56] Van Kempen, S. S. Kim, C. Tumescheit, M. Mirdita, J. Lee, C. L. M. Gilchrist, J. Söding, and M. Steinegger. Fast and accurate protein structure search with foldseek. Nature Biotechnology, 42(2):243–246, 2024.

[57] J. L. Watson, D. Juergens, N. R. Bennett, B. L. Trippe, J. Yim, H. E. Eisenach, W. Ahern, A. J. Borst, R. J. Ragotte, L. F. Milles, and et al. De novo design of protein structure and function with RFdiffusion. Nature, 620(7976):1089–1100, 2023.

[58] T. Widatalla, Z. Rollins, M.-T. Chen, A. Waight, and A. C. Cheng. Abprop: Language and graph deep learning for antibody property prediction. The 2023 ICML Workshop on Computational Biology, 2023.

[59] R. Zheng, S. Dou, S. Gao, Y. Hua, W. Shen, B. Wang, Y. Liu, S. Jin, Q. Liu, Y. Zhou, L. Xiong, L. Chen, Z. Xi, N. Xu, W. Lai, M. Zhu, C. Chang, Z. Yin, R. Weng, W. Cheng, H. Huang, T. Sun, H. Yan, T. Gui, Q. Zhang, X. Qiu, and X. Huang. Secrets of rlhf in large language models part i: Ppo, 2023.

